# Naked mole-rat and Damaraland mole-rat exhibit lower respiration in mitochondria, cellular and organismal levels

**DOI:** 10.1101/2021.08.05.455346

**Authors:** Kang Nian Yap, Hoi Shan Wong, Chidambaram Ramanathan, Cristina Aurora Rodriguez-Wagner, Michael D. Roberts, David A Freeman, Rochelle Buffenstein, Yufeng Zhang

## Abstract

Naked mole-rats (NMR) and Damaraland mole-rats (DMR) are the only two eusocial mammals known. Both species exhibit extraordinary longevity for their body size, high tolerance to hypoxia and oxidative stress and high reproductive output; these collectively defy the concept that all life-history traits should be negatively correlated. However, when life-history traits share similar underpinning physiological mechanisms, these may be positively associated with each other. Here, we propose that the bioenergetic properties of mole-rats share a potential common mechanism. We adopted a top-down perspective measuring the bioenergetic properties at the organismal, cellular, and molecular level in both species and the biological significance of these properties were compared with the same measures in Siberian hamsters and C57BL/6 mice, chosen for their similar body size to the mole-rat species. We found mole-rats shared several bioenergetic properties that differed from their comparator species, including low basal metabolic rates, a high dependence on glycolysis rather than on oxidative phosphorylation for ATP production, and low proton conductance across the mitochondrial inner membrane. These shared mole-rat features could be a result of evolutionary adaptation to tolerating variable oxygen atmospheres, in particular hypoxia, and may in turn be one of the molecular mechanisms underlying their extremely long lifespans.

## Introduction

Energy is the currency of life [1]. As the theoretical biologist Alfred Lotka famously stated, “In the struggle for existence, the advantage must go to those organisms whose energy-capturing devices are most efficient in directing available energies into channels favorable to the preservation of the species” [2]. Life-histories of animals should therefore reflect the allocation of metabolic energy to those traits that determine fitness [3–6]. Ironically, although metabolic energy is the fundamental currency of fitness, few life-history studies directly focus on bioenergetics [7].

A key premise of life-history theory posits that life-history traits are negatively associated with each other and are considered “trade-offs” [8]. The only known eusocial mammals, the naked mole-rats (NMR; *Heterocephalus glaber*) and the Damaraland mole-rats (DMR, *Fukomys damarensis*) appear to challenge this concept of life-history trade-offs [9, 10]. Both species exhibit extreme longevity on the basis of their body size and a delayed/retarded aging phenotype [11, 12]. Although less is known about the DMR age-related biology than that of the NMR [12, 13], breeders of both species show no menopause and demonstrate enormous reproductive outputs (60–140 times greater than other rodents [14]), yet are often the most long-lived individuals in their colonies [15, 16]. Such observations directly challenge the most prominent life-history trade-off, the cost of reproduction at the expense of somatic maintenance resulting in a negative association between reproductive output and longevity [17]. More impressively, besides their extreme longevity and high reproductive output, both these species also share other characteristics such as hypoxia tolerance [18] and tolerance of oxidative stress [19], features considered evolved traits in response to life below ground for millenia [20]. In spite of the similarities of these exceptional life history traits within this monophyletic clade [21], NMR and DMR exhibit fundamental differences in their evolutionary history: studies suggest they evolved eusociality independently [22, 23], and have divergent physiological responses to different stressors [23]. Apart from some studies examining metabolic rates and the energetics of thermoregulation [24–27], the bioenergetic properties of these mole-rats has been largely overlooked. Both species have basal metabolic rates (BMR) considerably lower than predicted allometrically [25]. A few studies using isolated mitochondria from both species have yielded equivocal and conflicting results [28–34]. Clearly, a comprehensive evaluation of bioenergetics of these two mole-rat species and their involvement in life-history traits are overdue.

Mode-of-life theory suggests that ecological traits, such as fossoriality, are positively correlated with lifespan [35]. Fossorial animals are protected from predation, airborne contagious agents and unfavorable climatic conditions and this affords protection from extrinsic mortality and contributes to a longer lifespan [14]. A subterranean lifestyle has to contend with reduced gas exchange through soil, leading to a marked decline in oxygen availability in deep burrows and nests where multiple animals rest together. This could also potentially induce bioenergetic adaptations that may further contribute to the prolonged lifespans of underground dwelling species. In the present study, we employed an integrative approach to concurrently measure bioenergetic properties of DMR and NMR at the organismal, cellular and organelle level. We also investigate the significance of these properties by comparing these findings with shorter-lived, above-ground dwelling laboratory rodents, such as the Siberian hamster (*Phodopus sungorus*) and the C57BL/6 mouse. We used the hamster and C57BL/6 mouse as comparisons since these species were housed in the same animal facility as DMR and NMR respectively. We believe this integrative approach will enable us to obtain synergistic feedback and valuable insights to understand the biological mechanisms governing and the aging process and longevity.

## Methods

### Animals

The husbandry of DMR and Siberian hamsters and the study procedures were conducted under University of Memphis IACUC permits (#0862; #0797). The husbandry of NMR and C57BL/6 mice and the study procedures were conducted at Calico Life Sciences [36]. All experimental procedures were approved by the Calico IACUC committee (B-2-2020). Please see the electronic supplementary material, methods for more information on the husbandary of these animals.

### Whole organism basal metabolic rate measurement

A total of ten young adult hamsters (4 months old; 5 male and 5 female), eleven young DMR (4-5 years old; 6 male and 5 female), nine middle-aged DMR (9-10 years old; 4 male and 5 female), and eight old DMR (16-20 years old; 4 male and 4 female) were used for whole organismal BMR measurements. Only non-breeding animals were used in the study. All BMR measurements were conducted using a standard flow-through respirometry system (Sable Systems International, Las Vegas, NV, USA) following to that described in Yap *et al.* [37] (electronic supplementary material, methods). BMR was measured between 9:00 and 17:00 at 30°C, which is within the thermoneutral zone for both DMR and hamster [38, 39]. BMR calculations were done based on the lowest averaged 3 min of oxygen consumption per measurement sequence after 4 h according to Lighton’s equations 10.6 and 10.7 [40] with ExpeData software, version 1.2.6 (Sable Systems International). Age-related changes in BMR have been previously undertaken for NMR using the identical methodology [41] and similarly considerable data is available for the BMR of mice [42].

### Primary fibroblast isolation and culture

Primary dermal fibroblast from NMR and C57BL/6 mice was obtained from Buffenstein lab as previous described [43]. NMR cells were initially propagated at 32 °C with 5% CO_2_ and 3% O_2_ before being acclimated to 37 °C overnight before cellular respiration measurements according to Swovick *et al.* [44] whereas fibroblasts of mice were isolated using identical procedures but maintained in incubators at 37 °C. Primary lung fibroblasts from DMR and hamsters and primary dermal fibroblasts were isolated from two three year old DMR and two four month old hamsters according to Seluanov *et al.* [45] and Zhang *et al.* [46].

### Cellular respiration measurements

Respiration in cells were measured using Seahorse XF96 Analyzers (Agilent, Santa Clara, CA) according to Wong *et al.* [47] (electronic supplementary material, methods). Rates of oxygen consumption and extracellular acidification are expressed relative to the protein content of the appropriate well. Rate of ATP production during baseline from glycolysis and oxidative phosphorylation were calculated according to manufacturer protocols [48].

### Mitochondria isolation

Lung (DMR and hamsters) and heart (NMR and C57BL/6 mice) tissues were immediately dissected out upon euthanasia. Lung mitochondria were isolated from lung tissue of DMR (n = 5; 4 years old) and hamster (n = 5; 4 month old) according to Spear *et al.* [49]. Heart mitochondria were isolated from hearts of NMR and C57BL/6J mice in ice-cold heart sucrose buffer (electronic supplementary material, methods). Regardless of species, the resultant supernatant was discarded, the final mitochondria pellets were suspended in ice-cold Mitochondrial Assay Solution and were kept at high concentration (~20 mg protein/mL) on ice until use according to Mookerjee *et al.* [50].

### Mitochondria respiration measurement

Body temperature varies for the various species used in this study: NMR (32-34 °C), DMR (~35.2°C), mouse (36.2 - 38 °C) and hamster (36.1 - 38 °C) [51]. In order to compare mitochondria as machineries, respiration chamber temperature should be standardized since temperature could influence mitochondrial respiration. Munro *et al.* [31] had previously shown the respiration of isolated mitochondria did not vary significantly between 30 °C and 37 °C for NMR but differed significantly for mouse. As a result, we used 37 °C as respiratory chamber temperature for all four species. Lung mitochondria (0.35 mg/ml) respiration was measured in MAS-1 at 37°C using high resolution respirometry (Oroboros O2k, Innsbruck, Austria). The rates of respiration of heart mitochondria isolated from NMR and C56BL/6J mice were assessed according to Rogers *et al.* [52] by Seahorse XFe96 extracellular flux analyzer (electronic supplementary material, methods).

### Western blot

Western blots were conducted on lung samples to analyze hypoxia-inducible factor 1-alpha (HIF-1α, GTX37356, GeneTex) according to Zhang *et al.* [53]. Each membrane was stained with Ponceau and was used as the loading and transfer control. A chemiluminescent system was used to visualize marked proteins (GE Healthcare Life Sciences, Pittsburgh, PA). Images were taken and analyzed with the ChemiDoc imaging system (Bio-rad, Hercules, CA).

### Statistical analyses

All statistical tests were carried out using IBM SPSS, version 26.0. BMR and body mass were log10 transformed prior to analyses. We first used general linear models (GLM) including the factor sex to compare BMR between DMR and hamster. However, adding or removing sex as covariant yielded similar results, so it was removed from statistical analyses. We tested the effect of species on BMR using GLM with BMR as dependent variable, species as main effect, and body mass as covariate [54]. To investigate if BMR differed between DMR from different age groups, we tested the effect of age group on BMR of DMR using GLM with BMR as dependent variable, age group as main effect, and body mass as covariate. *F*- and *t*- statistics and P values were reported. Student *t*-test was used for comparison in cellular and mitochondrial levels between NMR and DMR with their counterparts. We considered *P* < 0.05 as statistically significant.

## Results

### Whole organismal BMR

Our measurements of BMR in captive DMR are consistent with the reported BMR of this species captured in the wild [38] and in captivity [55]. BMR for hamsters were also consistent with previous studies [39] (Supplementary Table 1). Both DMR (*P* = 0.029) and hamsters (*P* = 0.046) demonstrated a statistically significant positive correlation between their body mass and BMR. We therefore included body mass as a covariant when performing comparisons of metabolic rates. We observed that DMR demonstrated a 50% lower BMR than hamsters (*F*_*1,*18_ = 4.55, *P* = 0.047; Figure 1 A,C), which aligned with predictions by Lovegrove [38]. We detected no significant differences between young, middle-aged and old DMR (*F*_2,24_ = 2.15, *P* = 0.138; Figure 1 B,D). Body mass also stayed constant between age groups in DMR (*F*_2,25_ = 0.99, *P* = 0.386; Supplementary Table 1). This finding is consistent with previous reports on NMR, where no age-related changes in body mass and BMR were found [41].

**Figure 1.**
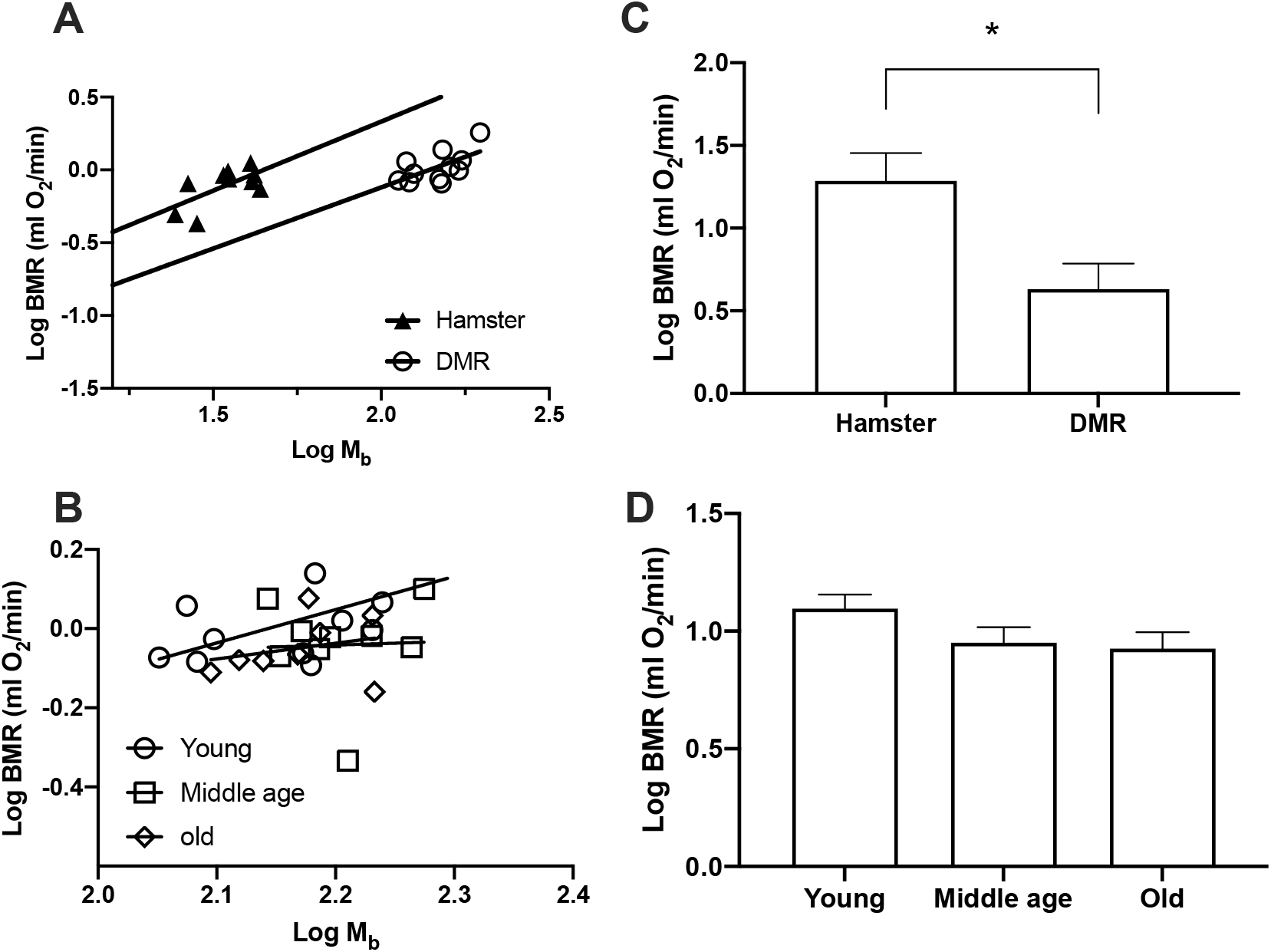
Basal metabolic rate (BMR) in Damaraland mole-rats (DMR) and Siberian hamsters. Least-squares linear regressions of log_10_-transformed BMR against log_10_ body mass (M_b_) for (A) DMR and hamsters, (B) young, middle age, and old DMR. Least-squares means (± s.e.m.) for each metabolic rate (C) DMR and hamsters, (D) young, middle age, and old DMR. * indicates *P* < 0.05

### Primary fibroblasts respiration

To better understand our observations, we move on to explore cellular bioenergetics of both of these species. We did so by measuring the rates of cellular respiration and the rates of glycolysis, the two main pathways to generate energy, using an extracellular flux analyzer. Primary lung fibroblasts of DMR showed significantly lower rates of basal (*t*_4_ = 5.47, *P* = 0.005; Figure 2 A,B) and FCCP-induced maximal (*t*_4_ = 7.61, P = 0.002; Figure 2 A,B) respiration when compared with hamster lung fibroblasts. Oligomycin-induced state 4O respiration, indicative of proton leak across mitochondrial inner membrane and the effectiveness of mitochondrial electron transport, were not different between DMR and hamsters (*P* = 0.60). Interestingly, basal extracellular acidification rate, which reflects basal glycolytic rates, were significantly higher in lung fibroblasts of DMR than of hamsters’ (*t*_4_ = 8.06, *P* = 0.001; Figure 2 C,D). The maximal capacity of glycolysis of these cells showed no differences between DMR and hamsters (*P* = 0.15). We also calculated the rates of ATP production based on the rates of respiration and glycolysis and observed no statistically significant differences of ATP production rates in lung fibroblasts of DMR and hamsters (*P* = 0.230; Figure 3A).

**Figure 2.**
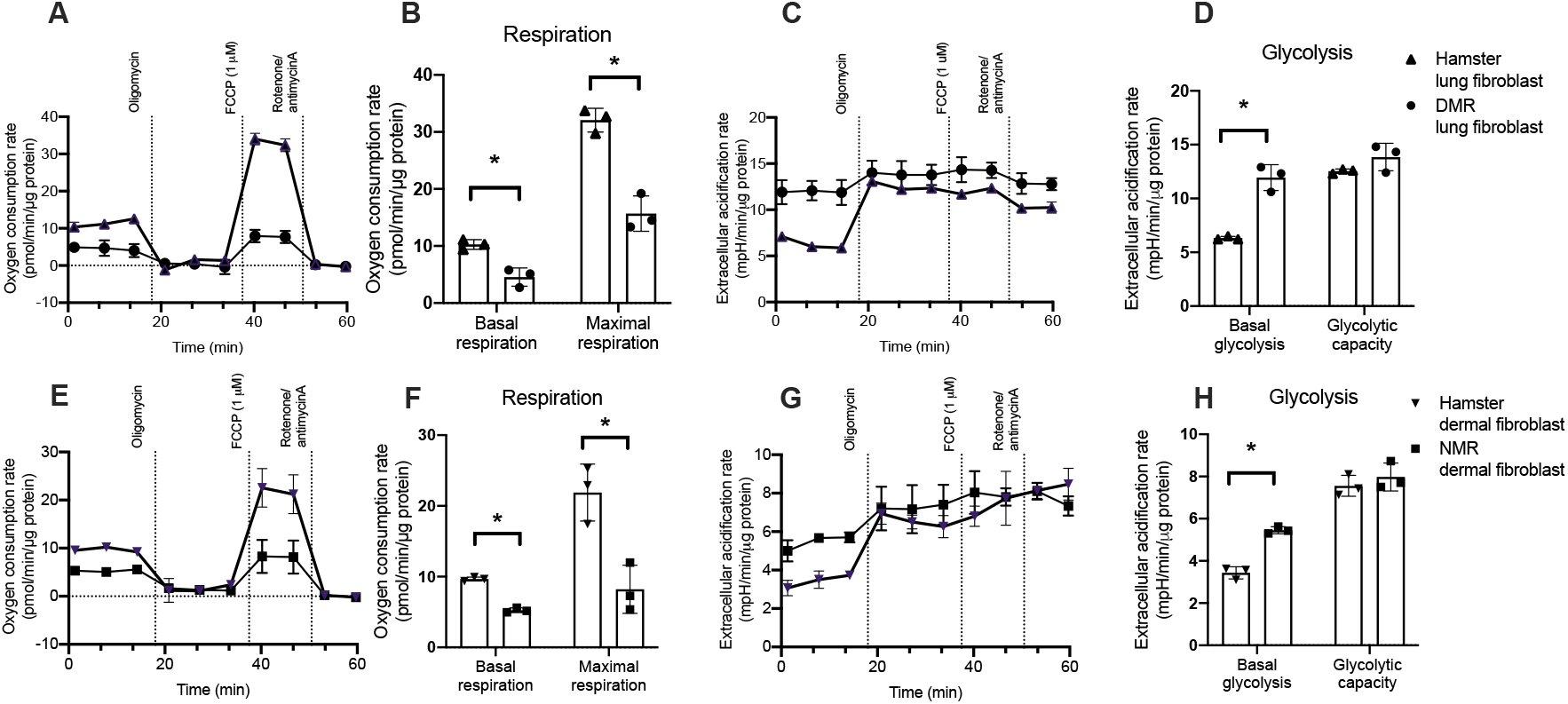
Respiration and glycolysis for (A-D) Damaraland mole-rat (DMR) and Siberian hamster lung fibroblast, and (E-H) naked mole-rat (NMR) and Siberian hamster dermal fibroblast. Seahorse traces for oxygen consumption (A, E) and extracellular acidification (glycolysis; C,G) rate in fibroblasts. B, F: calculated basal and maximal respiration. Basal and Maximal respirations for hamster lung fibroblast were indicated by shaded area in A as an example. D, H: calculated basal glycolysis and glycolytic capacity. Basal glycolysis and glycolytic capacity for hamster lung fibroblast were indicated by shaded area in C as an example. N = 3 independent experiments, Data are shown as means ± s.e.m., * indicates P < 0.05.

**Figure 3.**
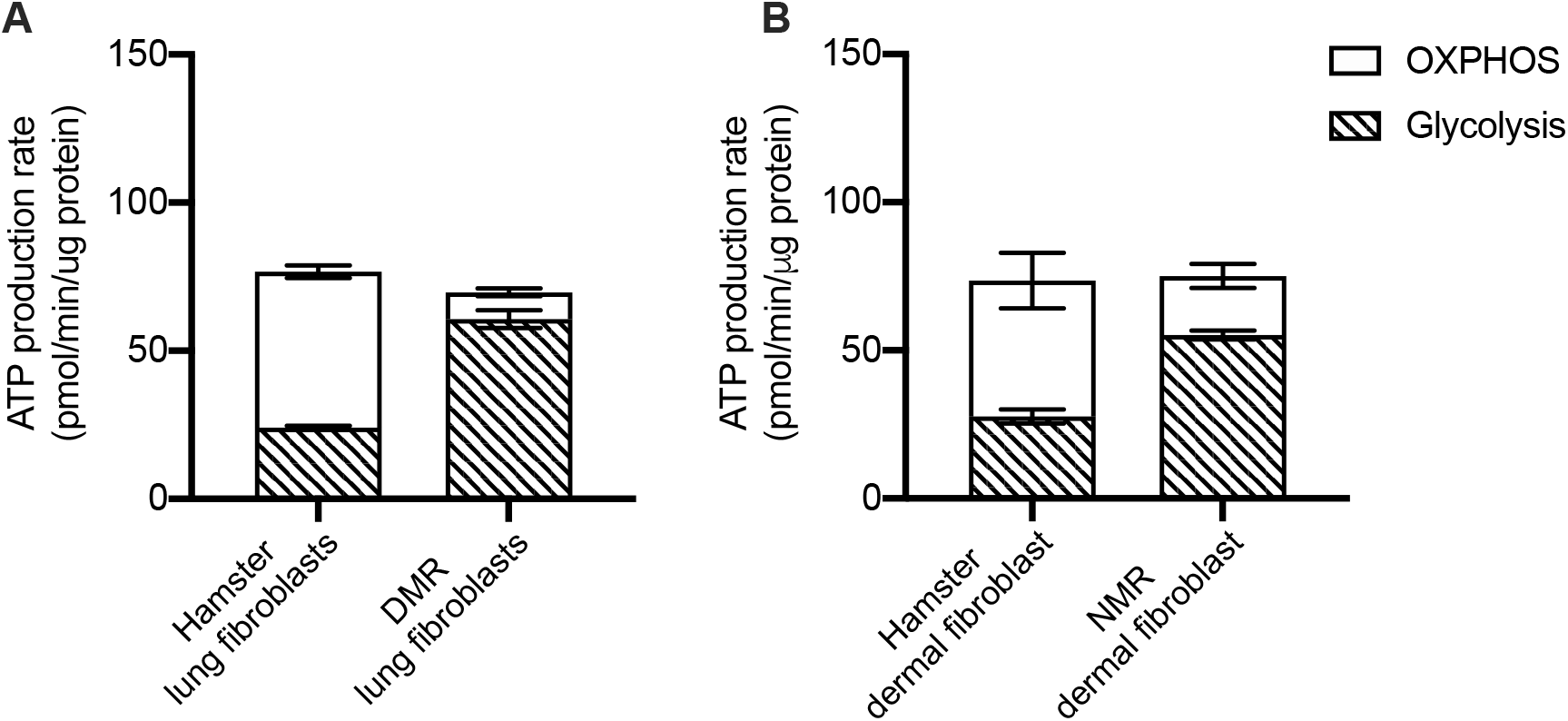
ATP production rates during baseline from oxidative phosphorylation (OXPHOS) and glycolysis in (A) Damaraland mole-rat (DMR) and hamster lung fibroblast, (B) naked mole-rat (NMR) and hamster dermal fibroblast. N = 3 independent experiments, Data are shown as means ± s.e.m., * indicates P < 0.05

We extended our investigation to also cover primary dermal fibroblasts of NMR. Intriguingly, similar patterns were observed between the comparisons between NMR and hamster dermal fibroblasts. Dermal fibroblasts of NMR demonstrated significantly lower rates of basal (*t*_4_ = 17.49, *P* < 0.001; Figure 2 E,F) and maximal respiration (*t*_4_ = 4.49, *P* = 0.011; Figure 2 E,F), whereas the rates of state 4 respiration were not different (*t*_4_ = 0.31, *P* = 0.77). While the maximal glycolytic rates were similar between dermal fibroblasts of NMR and hamster (*P* = 0.43), we again observed a significantly higher rates of basal glycolytic rates in NMR dermal fibroblasts (*t*_4_ = 10.44, *P* < 0.001; Figure 2 G,H). No differences in ATP production rates were detected between dermal fibroblasts of NMR and hamster (*P* = 0.904; Figure 3B).

### Mitochondrial respiration

Investigation using isolated mitochondria allows us to better understand the bioenergetic machinery employed during respiration and may yield some explanations for their disparate bioenergetic characteristics observed at either the cellular or organismal level (Supplementary Figure 1,2).

The rates of mitochondrial respiration were examined using both complex I-linked and complex II-linked substrates. In general, despite isolating mitochondria from different tissues (lungs: hamster and DMR; hearts: NMR and mice) the isolated mitochondria of DMR and NMR showed similar patterns that differed from those of hamsters and mice respectively. While the rates of Complex I-driven mitochondrial state 3_ADP_ respiration were found to be similar when comparing the mole-rats to their laboratory counterparts (DMR vs. hamsters: *P* > 0.288; Figure 4A; NMR vs. mice: *t*_14_ = 2.06, *P* = 0.058; Figure 4B), the rates of state 4o respiration were found to be significantly lower for both DMR and NMR than those of hamsters or mice (DMR vs. hamsters: *t*_8_ = 4.21, *P* = 0.003; Figure 4A; NMR vs. mice: *t*_14_ = 2.96, *P* = 0.010; Figure 4B). Collectively, this gives rise to mole-rats having significantly higher respiratory control ratios (RCR), a reliable indicator of mitochondrial bioenergetic integrity (*P* < 0.01; Supplementary Figure 3B).

**Figure 4.**
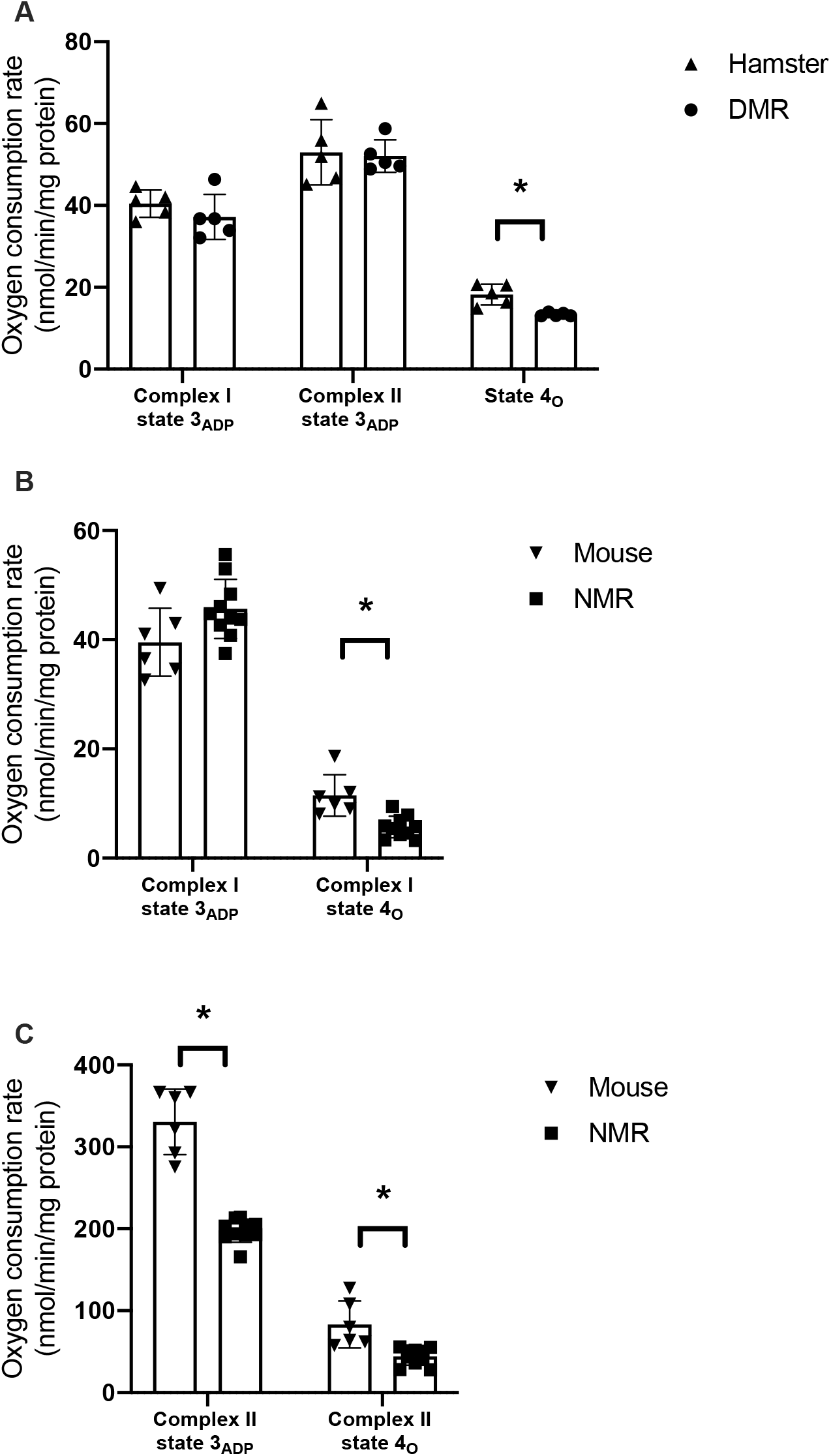
Respiration rate for (A) isolated lung mitochondria from Damaraland mole-rat (DMR; N = 5) and Siberian hamster (N = 5) measured using Oroboros O2k, (B) isolated heart mitochondria from naked mole-rat (NMR; N = 10) and C57BL/6 mouse (N = 6) measured using Seahorse XF96 Analyzers. Data are shown as means ± s.e.m., * indicates P < 0.05.

We also observed some degree of species-specific characteristics. When comparing DMR and hamsters, no differences were observed for Complex II-driven state 3_ADP_ respiration in lung mitochondria. However, we observed a significant decrease in the rate of Complex II-driven state 3_ADP_ respiration in heart mitochondria of NMR when compared to that of mice. The slower Complex II-driven state 3_ADP_ respiration observed here were consistent with previous findings [28, 29]. Such low levels of respiration by complex II driven substrates in NMR mitochondria were tied to decreased ROS production rate from complex I [28, 56].

## Discussion

Here we report that both DMR and NMR exhibited similar bioenergetic properties at the organismal, cellular and organelle levels compared to their above-ground dwelling, shorter lived but similar sized laboratory counterparts. Most notably both species have lower BMR than above-ground dwelling animals, cellular respiration is similarly lower while mitochondrial RCR is also higher. These data suggest convergent evolutionary processes most likely have arisen in response to social living under the variable oxygen atmospheres encountered below ground.

BMR of the two mole-rat species have been estimated to be about 30% - 60% lower than that predicted by body mass [12, 25, 27, 38, 41, 51]. The comparison between these two mole-rats species and other subterranean rodents, other partial fossorial mole-rat species, also shows a reduced BMR by more than 29% [38]. The observed lower BMR might be explained by their fully fossorial lifestyle and eusocial colony organization which likely results in repeated exposure to severe hypoxia [18]. Interestingly, while age-related declines in BMR in disease-free individuals are extensively documented feature in human and laboratory rodent species [57, 58], we observed no detectable age-related declines in body mass (Mb) and BMR in DMR or the NMR [41]. This likely is indicative of well-maintained body composition, most notably lean mass, tissue function, and bioenergetic properties with increasing age [59].

BMR was calculated through oxygen consumption measurements. To understand the mechanisms underlying lower BMR in mole-rats, we extended our investigation to measure cellular respiration using primary fibroblasts. We found that fibroblasts from both mole-rat species exhibit lower rates of basal and maximal cellular respiration than that of the hamster, with associated elevations in basal glycolytic rates. We further calculated the rates of total ATP production based on the rates of basal cellular respiration and glycolysis. We observed that the rates of ATP production of primary fibroblasts isolated from the two mole-rat species were comparable to those of hamster in spite of reduced basal cellular respiration rates in the two mole-rat species (Figure 3). This intriguing finding may be explained by the compensatory mechanism between mitochondrial and glycolytic ATP production during which either one of the pathways can be upregulated upon the slowdown of the other. Our observations here suggest a higher reliance of mole-rat fibroblasts on glycolysis instead of oxidative phosphorylation (OXPHOS) for ATP production. Although Swovick *et al.* [60] similarly showed lower basal respiration levels in NMR compared to mice; inconsistent with our findings, that study reported lower rates of basal glycolysis with concomitant lower total ATP production compared to that of mouse cells. The exact reason for this interstudy discrepancy is still unclear. However, Swovick *et al.* [60] conducted their measurements in confluent, quiescent cells, whereas our measurements were performed in normal growing cells. It is possible that normal growing cells may have a higher ATP demand which is met by an increased basal glycolysis in mole-rat cells. Despite this discrepancy, consistent with our observation Swovick *et al.* [60] also observed a higher percentage of ATP production from glycolysis than OXPHOS in NMR compared to mouse cells, in keeping with the Warburg effect [61]. This preference for glycolysis over OXPHOS reportedly elicits a broad range of beneficial health span and/or lifespan consequences [61]. For example, pharmacological and genetic manipulations that moderately inhibit mitochondrial respiration, such as metformin, TPP-thiazole, reportedly extend lifespan [62–64] by shifting ATP production from mitochondrial OXPHOS to glycolysis. The observed similar bioenergetic pattern favoring ATP generation through glycolysis is beneficial under hypoxic conditions that limit oxygen availability for OXPHOS and may indirectly also contribute to their extremely long lifespan.

Isolated mitochondria from both mole-rat species had lower state 4_O_ respiration. State 4_O_ respiration is used as an indirect measure of the proton conductance across the mitochondrial inner membrane and serves as an indicator of mitochondrial coupling. Increased mitochondrial coupling is suggestive of enhanced efficiency of mitochondrial ATP production per unit of oxygen consumed, findings consistent with previous reports on isolated mitochondria from liver, heart and skeletal muscle of NMR [29–31]. The mitochondria isolated from the two mole-rat species in this current study, were much more coupled than those of hamsters and mice.

Lau *et al.* [29] have observed a lower state 4_O_ respiration and a concomitant increase in mitochondrial membrane potential (ΔΨ) in NMR heart mitochondria, which indicated that mitochondria were hyperpolarized during state 4_O_ respiration. Consequently, the observed lower mitochondrial proton conductance may be due to differences in either trans-membrane transporters (such as uncoupling proteins (UCPs), adenine nucleotide transport) or inner membrane phospholipid composition [65]. Upregulation in NMR uncoupling proteins [66] or a lower proportion of the polyunsaturated fatty acid docosahexaenoic acid in the mitochondrial inner membrane [67] would reduce proton conductance across the inner membrane and therefore contribute to the observed lower state 4_O_ respiration measured in NMR in both this and Lau’s study. It is worth noting that some previous studies had reported contradictory results [32–34]; those studies suggest that NMR mitochondria are more uncoupled than other laboratory rodents, as indicated by their findings of a lower ΔΨ during state 4 respiration when compared to mice. Their observations support the “Uncoupling to Survive” hypothesis which proposes that mitochondrial uncoupling would result in decreased reactive oxygen species production, reducing oxidative stress with concomitant effects of increased longevity [68]. Further studies are needed to determine why there is so much interstudy variability and if this rather reflects disparate acclimation or other laboratory-specific experimental conditions. Inevitably, uncoupled mitochondria would produce more heat than those that are well-coupled, facilitating better maintenance of body temperature. Given that NMR are extremely thermolabile, unable to defend body temperature at ambient temperatures even a few degrees below thermoneutrality [24, 25], we believe our observation is more consistent with their physiology.

One of the strengths of the present study is the use of an integrative approach to concurrently measure respiration at the organismal, cellular and organelle levels. Such approach enables synergistic feedback of insights from observations at all three levels of organization, and is more informative than conducting such studies at a single biological level. In this context, we observed a mismatch between results from isolated mitochondria and the findings obtained from cellular respiration in present study. Most notably, we observed lower rates of state 4 and similar rates of state 3 respiration in isolated mitochondria, but lower rates of state 3 and similar rates of state 4 were seen in cells from mole-rats compared to their counterparts. In isolated mitochondria assays, excessive substrates were provided during state 3 respiration. As a result, the differences between state 3 respiration on mitochondria and cells indicated the substrates availability might be different between mole-rats fibroblasts compared to their counterparts. The activation of UCPs and other factors might influence mitochondrial proton conductance in cells and could explain the differences between state 4 respiration in mitochondria and cells [69]. These results suggest that mitochondria could behave differently in cells compared to when they are isolated and supplied with unlimited substrates. Consequently, we need to be cautious when extrapolating data solely obtained from isolated mitochondria to explain physiological phenomena.

It has also been hypothesized that mitochondria basal proton leak (represented by state 4 respiration) significantly contributes to BMR *in vivo* [70]. In this study we observed significantly lower mitochondrial basal proton leak, cellular respiration and BMR. However, it is important to note that other factors may also contribute to the magnitude of species differences in BMR; mitochondrial basal proton leak was 39% - 43% lower depending on both the substrates and species, basal respiration in isolated primary dermal fibroblast cultures were reduced by 45% - 55% although rates of ATP production were similar, and BMR was 50% - 60% lower for two mole-rat species compared to their counterparts [71].

Our study employed both NMR and DMR to evaluate bioenergetic profiles of subterranean species. Although these two species have similar life-history traits, they are fundamentally different in their evolutionary history. Phenotypic convergence could be the result of similar adaptation to shared environments [72]. Consequently, convergent evolution can serve as a valuable proxy for repeated evolutionary experiments in nature. Moreover, understanding how convergent traits evolve, especially at different biological levels, has the potential to inform general rules about adaptation [73]. Mode-of-life theory suggests that ecological traits, such as fossoriality, are positively correlated with lifespan [35]. Strictly subterranean animals can escape predation and unfavorable above-ground conditions, these reductions in extrinsic mortality also may contribute to longer lifespan [14]. But in the physiological context, it is possible that the adaptations for life below ground where low oxygen atmospheres may be common contribute to the unique bioenergetic properties shared by these two mole-rat species and as a byproduct thereof also could contribute to their longevity. Adaptation to hypoxic environment includes decline in BMR at the organismal level [74], higher reliance on glycolysis than OXPHOS for ATP production at the cellular level [75], and decreased mitochondrial proton conductance at the organelle level [76] traits observed in both mole-rat species.

Although the exact physiological pathways underlying these bioenergetic adaptations are unknown, a top candidate may be hypoxia-inducible factor 1 (HIF-1). HIF-1, or its alpha subunit (HIF-1α). HIF-1α is considered as the master transcriptional regulator of cellular response to hypoxia [77]. In most mammals, HIF-1α is only activated during hypoxia and is constantly being degraded during normoxia. In NMR, genomic analysis indicated several mutations can prevent HIF-1α from degradation during normoxia, which would chronically upregulate HIF-1α mediated signaling pathways [66]. In DMR, HIF-1α protein level in lungs was much higher than in hamsters and similarly NMR constitutively have higher HIF-1a than observed in mice (Supplementary Figure 4). Interestingly, HIF-1 activation could also delay cellular senescence [78], and extend lifespan in many different model species [79, 80]. It is likely that adaptation to the hypoxic environment upregulate HIF-1 regulated pathways resulted in these unique bioenergetic properties. The persistent upregulation of HIF-1 pathways and the unique bioenergetics may elicit beneficial effects to promote the extremely long lifespan of these mole-rat species. Such hypothesis is in line with recent studies that indicated that instead of trade-offs, different life-history traits could be positively associated with each other, when these traits share similar underpinning physiological mechanisms [17, 53].

In summary, our study has shown similar bioenergetic properties shared by NMR and DMR at the organismal, cellular, and organelle levels. These bioenergetic characteristics, such as lower but relatively constant BMR throughout the life, dependence on glycolysis rather than mitochondrial OXPHOS for ATP production in cells, lower basal proton conductance through mitochondria inner membrane, could be resulted from adaptation to hypoxic environments. These metabolic adaptations, on the other hand, might be beneficial for other life-history traits such as longevity. Future studies directly testing these underlying mechanisms, together with studies that involve other fossorial or semi-fossorial rodents would provide important insights for changes in metabolism in hypoxic environments, and its effects on aging. More importantly, these studies can reveal mechanistically how different life-history traits interact and influence each other.

## Acknowledgement

We would like to thank the vivarium staff from both University of Memphis and Calico Life Sciences for their excellent animal care. We also thank Matthew Butawan, Megan Smith, Mary McMahon, and Wendy Craft for their technical support. Y.Z. was supported by US National Science Foundation grant IOS – 2037735, while Calico Life Sciences funded work undertaken there.

## Author Contributions

K-N.Y., H-S.W., D.A.F., R.B., and Y.Z. contributed to conceptualization, methodology, investigation, data curation, visualization, supervision, writing and editing. C.R., C.A.R., and M.D.R., contributed to investigation and editing.

