## Supplementary Materials for "Naked mole-rat and Damaraland mole-rat exhibit lower respiration in mitochondria, cellular and organismal levels"

Supplementary Methods:

*Animals*

Damaraland mole-rat (DMR) used in this study came from colonies housed at the University of Memphis (originally provided by Dr. Bruce Goldman at the University of Connecticut); their progenitors of which were collected in Namibia by Jennifer Jarvis. Their diet consisted of *ad libitum* sweet potatoes supplemented with dry rodent pellets (Harlan 2019, 19% protein diet). Each individual colony was housed within a complex constructed of two differently sized cages (60 × 40 × 20 cm^3^ and 48 × 25 × 20 cm^3^) connected by varying lengths of extruded polycarbonate tubing to roughly simulate natural burrow architecture. The number of cages and lengths of tube were dependent upon the sizes of the colonies and all cage contained a 1:1 mixture of corncob and pine bedding. DMR were maintained on a 12-hour dark-light cycle and were provided food *ad libitum*. Siberian hamsters used in this study also came from colonies housed at the University of Memphis (originally supplied by Irving Zucker at the University of California, Berkeley) in polypropylene cages (29 × 18 × 13 cm^3^) in 16 L:8 D at 22 ± 1 °C. Animals had access to food (8640 Teklad 22/5 Rodent Diet, Teklad Diets, Madison, WI) and water *ad libitum*.

C57BL/6 mice were purchased from the Jackson Laboratories (Bar Harbor, ME) and maintained in the vivarium for at least 2 weeks prior to use whereas the NMR used here were part of the well-characterized Buffenstein colony and were housed under well-defined housing conditions. Both species were maintained on a 12-hour dark-light cycle and were provided with food *ad libitum*. While NMR obtained all their water and nutrient requirements from the fresh fruits and vegetables that they were fed, mice were provided water *ad libitum* in addition to standard mouse chow.

*Whole organism basal metabolic rate measurement*

During measurements, animals were placed in one in each of the eight metabolic chambers (2.5-litre plastic canister, Nalgene). The system was checked for leaks before each round of metabolic rate measurement. All metabolic chambers were placed in an incubator (PELT-5 Peltier effect temperature-controlled portable cabinet, Sable Systems International) maintained at 30°C for the entire duration of BMR measurements. Each metabolic chamber continuously received ∼550 ml min^−1^ of dry air (using drierite as scrubber). Each of the seven metabolic chambers containing an animal and an empty chamber sampling baseline ambient air were sampled for 10 min by a multiplexer (RM8, Sable Systems International) every 80 min, allowing a minimum of 50 min of recording per chamber for 8 h. Animals were allowed to settle down in the chamber for at least 4 h to achieve post absorptive stages. Animals were weighed immediately before and after measurements and the average of the two masses was used in BMR analysis.

*Cellular respiration measurements*

Cells were seeded at a final density of 10,000 cells/well into an XFe96 cell culture microplate. After cell attachment, cells were washed with pre-warmed Krebs Ringer Modified Buffer (KRB: 35 mM NaCl, 5 mM KCl, 1 mM MgSO_4_, 0.4 mM K_2_HPO_4_,20m MHEPES and 5.5 mM glucose, pH 7.4 at 37 °C, supplemented with 0.1%w/v bovine serum albumin (KRB-BSA)) twice followed by a 30-min incubation with KRB-BSA in an air incubator at 37 °C. The rates of cellular respiration were then measured using Seahorse XF Cell Mito Stress Test Kit (Cat. No. 103015-100 from Agilent) according to corresponding manufacturer protocols. The maximum capacity of mitochondrial respiration was defined as the maximum rate induced by carbonyl cyanide p- trifluoromethoxyphenylhydrazone (FCCP, 1 μM). Following the assay, protein concentration in each well was measured by a BCA assay according to the manufacturer's instructions.

*Mitochondria isolation*

Lung tissues were excised and minced in 10 mL isolation buffer (250 mM sucrose, 2 mM EDTA, 5 mM Tris-HCI, and 0.5% BSA, pH 7.4). Minced tissues were then homogenized in a Potter-Elvehjem PTFE pestle and glass tube. The resulting homogenate was centrifuged at 500 × *g* for 10 min at 4°C, and the supernatant filtered through cheesecloth underwent another round of centrifugation at 10,000 x g for 10 min.

To isolate heart mitochondria from NMR and C57BL/6J mice, hearts were excised and were kept in 4 mL of ice-cold heart sucrose buffer (HSB: 320 mM sucrose, 1 mM EDTA, 50 mM Tris-HCl, pH 7.2) prior to homogenization. Hearts were homogenized in 4 mL of ice-cold HSB using a motor-driven Teflon in glass homogenizer. Homogenate was centrifuged at 500 × *g* at 4 ℃ for 10 min to remove tissue debris and nuclei. Supernatant was transferred into a clean centrifuge tube and was centrifuged at 10,000 × *g* at 4 ℃ for 10 min. The pellets were then washed by ice-cold HSB followed by another round of centrifugation at 10,000 × *g* at 4 ℃ for 10 min.

Regardless of species, the resulting supernatant was discarded and the final mitochondria pellet was suspended in Mitochondrial Assay Solution (MAS-1: 2 mM HEPES, 10 mM KH_2_PO_4_, 1 mM EGTA, 70 mM sucrose, 220 mM mannitol, 5 mM MgCl_2_, 0.2% w/v fatty acid-free BSA, pH 7.4).

*Mitochondria respiration measurement*

Lung mitochondria (0.35 mg/ml) respiration was measured in MAS-1 at 37°C using high resolution respirometry (Oroboros O2k, Innsbruck, Austria). Mitochondrial respiration was measured using both complex I (10 mM pyruvate, 10 mM glutamate) and complex II (10 mM succinate, 2 µM rotenone) substrates. State 3_ADP_ (maximal) respiration was induced by addition of 5 mM ADP, and state 4_O_ (state 4_Oligomycin_; idling) respiration was induced by addition of 2 µg/mL oligomycin. Respiratory control ratio (RCR) was calculated by dividing state 3 _ADP_ by state 4_O_ respiration.

The rates of respiration of heart mitochondria isolated from NMR and C56BL/6J mice were assessed by Seahorse XFe96 extracellular flux analyzer. To support Complex I-linked respiration, mitochondria were provided with 5 mM L-glutamate plus L-malate as respiratory substrates. To support Complex II-linked respiration, 5 mM succinate plus 4 uM rotenone were provided instead. The rates of respiration at basal state 2 (substrate only), phosphorylating state 3_ADP_ (5 mM ADP), non-phosphorylating state 4o (1 μg/mL oligomycin) and uncoupled respiration (4 μM FCCP) were measured in plate-attached mitochondria. The amount of mitochondria in each well was optimized for specific reactions to achieve optimal rates of respiration: 5 μg of mitochondrial protein when glutamate plus malate were respiratory substrates and 2 μg of mitochondrial protein when succinate plus rotenone were used. Respiratory substrates were added just before the assay plate was loaded into the instrument. Heart mitochondria of each of the animals were tested as quadruplicate on each plate. Measurements were performed at 37 °C in MAS-1 buffer containing 0.3% (w/v) BSA. Rates of mitochondrial respiration were expressed as nmol O_2_/min/mg protein.

Supplementary Table 1.

| Species | Age | Body mass | BMR (ml O_2_/min) |
| --- | --- | --- | --- |
| Siberian hamster, Phodopus sungorus | 4 month old | 35.10 ± 2.18 | 0.79 ± 0.08 |
| Damaraland mole-rat, Fukomys damarensis | 4 years old | 148.30 ± 8.02 | 1.16 ± 0.16 |
| Damaraland mole-rat, Fukomys damarensis | 8-10 years old | 160.48 ± 5.77 | 0.88 ± 0.07 |
| Damaraland mole-rat, Fukomys damarensis | 16-20 years old | 148.28 ± 5.96 | 0.9 ± 0.06 |

Supplementary Figure 1. Representative trace for Damaraland mole-rat and Siberian hamster lung mitochondria respiration using Oroboros O2k.

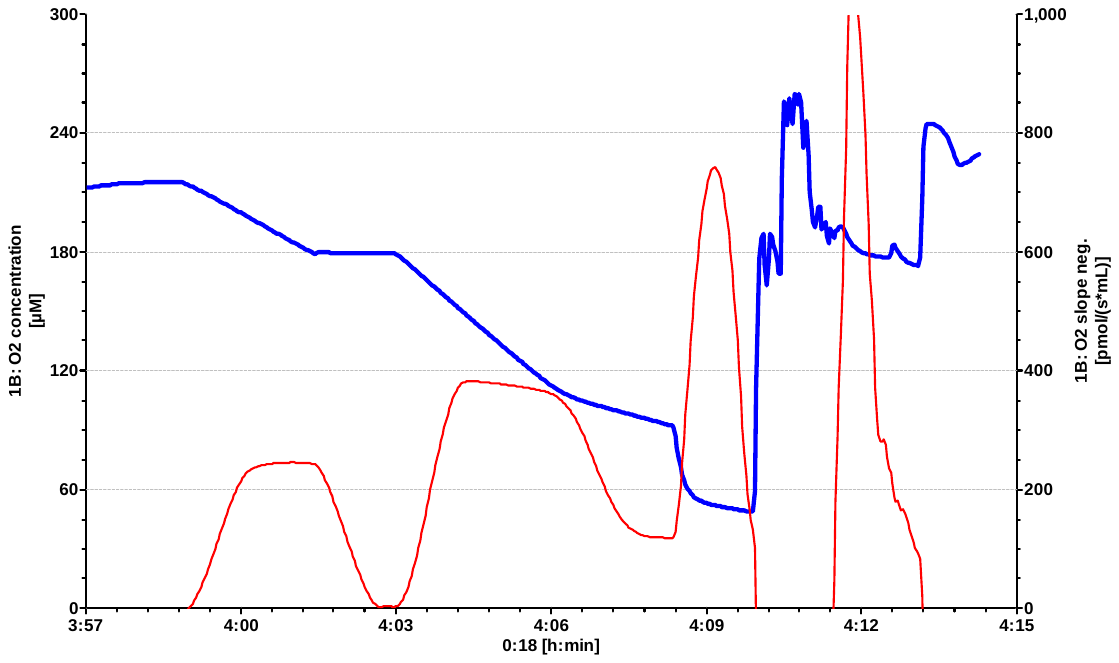

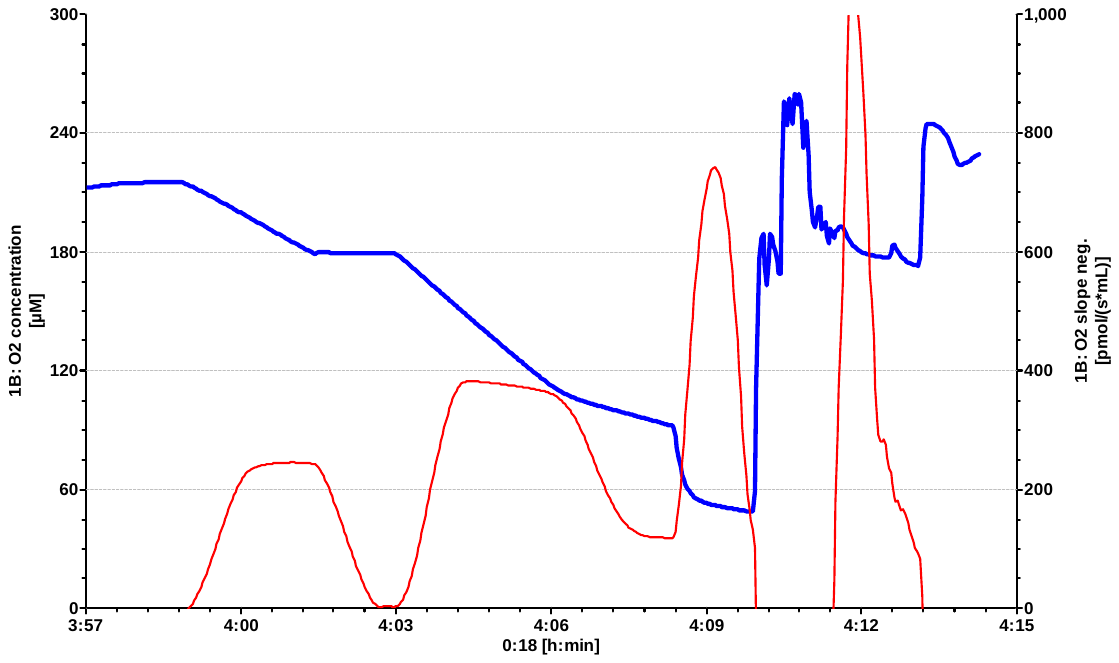

**Complex I**

**State 3**

**Complex II**

**State 3**

**State 4**

Mito + Complex I substrates

ADP

Rotenone

Complex II substrates + ADP

Oligomycin

**O_2_ Concentration (µM)**

**O_2_ Consumption (µmol/(s*mL))**

Supplementary Figure 2. Representative trace for naked mole-rat and C57BL/6 mice heart mitochondria respiration using Seahorse XFe96 extracellular flux analyzer. Complex I substrates: L-glutamate and L-malate; Complex II substrates: succinate and rotenone.

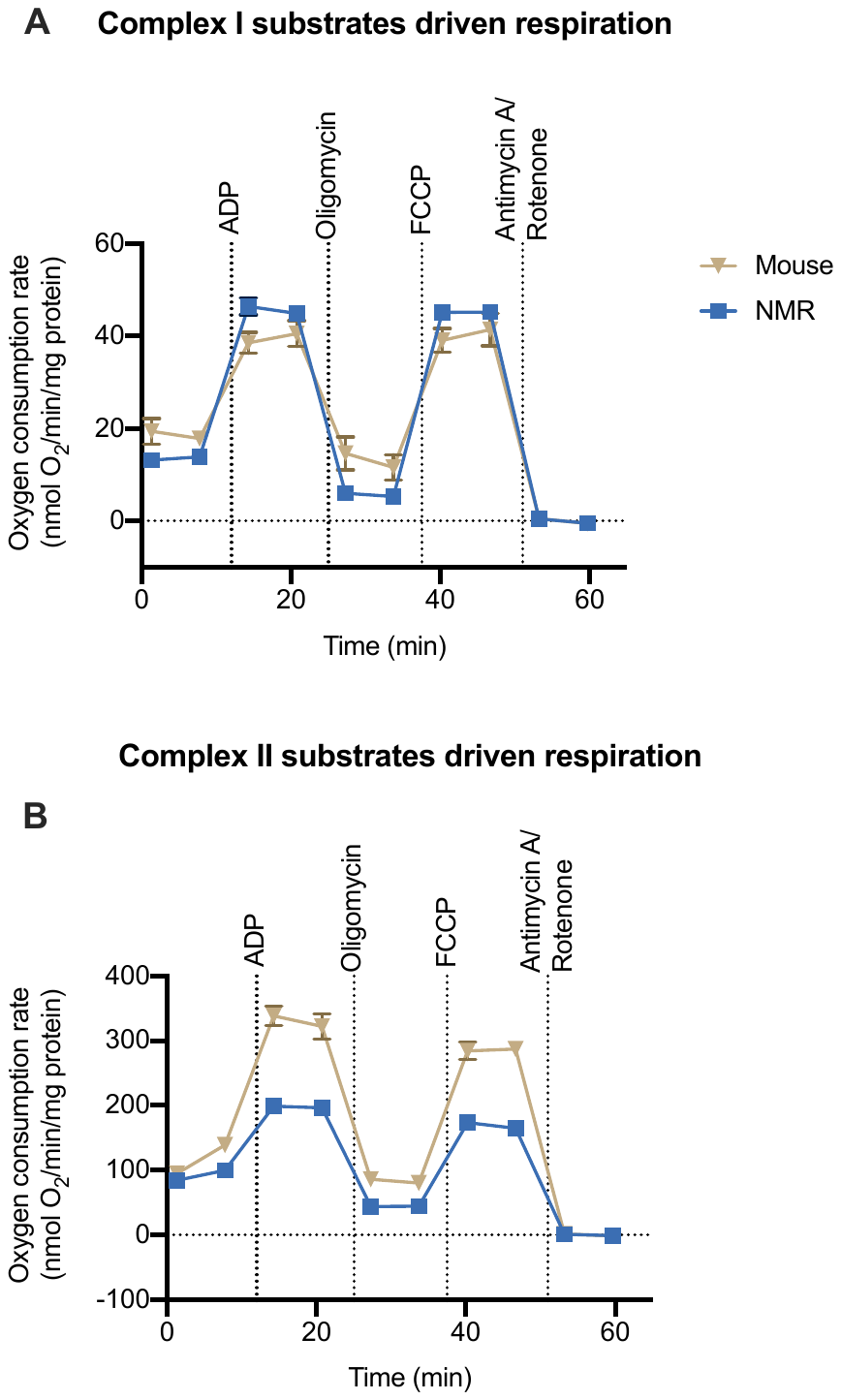

Supplementary Figure 3. Respiratory control ratio (RCR: State 3/ State 4) of (A) isolated lung mitochondria in Siberian hamster (N = 5) and Damaraland mole-rat (DMR; N = 5); (B) isolated heart mitochondria in C57BL/6 mice (N = 6) and naked mole-rat (NMR; N = 10). * indicates *P* < 0.05

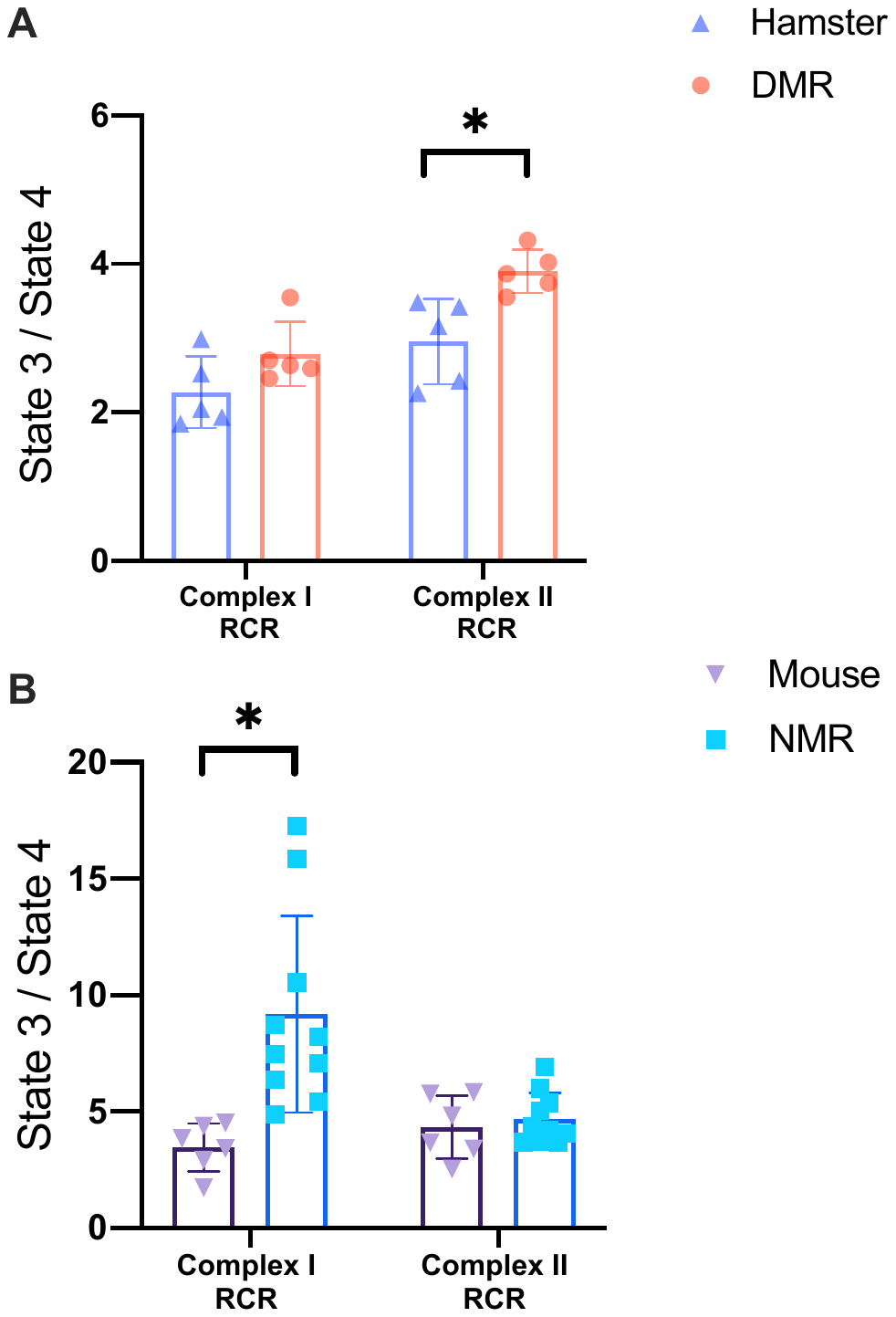

Supplementary Figure 4. Hypoxia inducible factor 1 aplha (HIF-1α) protein level measured by western blot in Damaraland mole-rat (DMR) and hamster lung tissues. * indicates P < 0.05

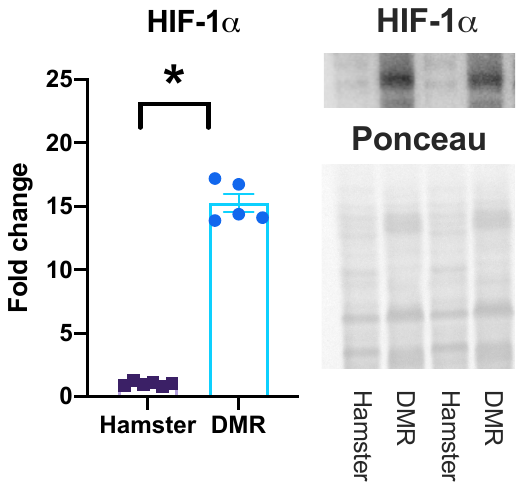
